# Microbial Communities in Cave Waters Across Karst Regions of Virginia

**DOI:** 10.64898/2026.08.09.738996

**Authors:** Riley S. Drake, Katarina Kosič Ficco, Thomas E. Malabad, William Orndorff

## Abstract

Karst groundwater supplies in Virginia are relied on to varying degrees for domestic, agricultural, and municipal water supplies. Further, Virginian caves harbor an estimated 200 endemic invertebrate species. The microbial occupants of Virginia’s karst aquifers are largely undescribed; characterizing them promises to inform both the scientific description of these systems and the management of a critical water resource. Karst aquifers are heterogeneous, and much of the water moving through them cannot be reached directly; we profiled cave waters both because cave passages offer direct access to active groundwater and because cave water specifically is relied upon by endemic invertebrate species living in caves. Using 16S rRNA sequencing, we characterized aquatic microbial communities in eight Virginia caves, across Virginia’s four major karst regions. We identified 3,899 unique amplicon sequence variants (ASVs) and found that caves hosted diverse microbial assemblages that differed markedly among sampled sites. These baseline data provide a starting point for future work to understand how seasonal cycles, weather events, and surface disturbances affect the microbial communities present in cave waters and the cave-endemic invertebrates that depend on these waters.

## Introduction

The faulted and folded geology of Virginia produces hydrologically isolated karst groundwater systems (1–3) that harbor an estimated 200 endemic, cave-limited invertebrate species (4–6) and are especially vulnerable to surface impacts and the associated loss of endemic species (7). These groundwater systems are part of a broader karst aquifer network that is relied upon, to varying degrees, for domestic, agricultural, commercial, and municipal water supplies (8, 9). Characterizing the microbial occupants of these aquifers therefore has considerable potential to inform both their scientific description and the management of these critical resources.

Karst aquifers are heterogeneous (10), and much of the water moving through them is inaccessible to direct observation. We profiled cave waters both because cave passages offer access to active groundwater and because cave water specifically is relied upon by endemic invertebrate species living in caves. Biological conservation of a cave system or karst area typically begins with an inventory of the species present (11). Biological inventory, for example of cave invertebrates, generally requires multiple visits before the full diversity of organisms present can be reliably estimated (12–16). Once resident taxa and their ranges have been established, the dynamics, vulnerability, and stability of the environment can be assessed through monitoring—monitoring involves repeated measurements, usually of one or more taxa of interest, made across time (17, 18). High-throughput approaches such as sequencing microbial or environmental DNA in cave water can be carried out during a single visit and offer information complementary to a traditional biological inventory (19, 20). We anticipate that microbial inventory of cave waters will eventually function much as other forms of biological inventory do, providing a starting point for studies aimed at monitoring how seasonal cycles, irregular weather, and other surface disturbances affect the microbial communities present in cave waters, which may also impact the stygobiontic invertebrates that depend on cave waters.

As an initial survey of the microbiology of cave waters across Virginia, we characterized the aquatic microbial communities of nine water samples collected from eight caves that span different carbonate units and karst regions across Virginia. We used full-length (V1–V9) 16S rRNA gene sequencing on the PacBio platform, which improves taxonomic resolution relative to short-read approaches (21), to describe community composition and diversity across these sites.

## Materials and Methods

### Sample Site Details

Nine cave water samples were collected from eight caves between April and November 2022 (Table 3). Sites spanned carbonate units ranging from Cambrian–Ordovician to Mississippian in age and were distributed across the four major karst regions of Virginia (Fig. 1). We detail the hydrogeological setting of each sample as metadata (Table 3): most samples were vadose cave streams, captured at either the input or the discharge end of the cave system, while the Madisons Saltpeter sample was collected from a small phreatic pool. Two samples (Omega Main Stream and Omega Tributary) were collected from separate streams at comparable depth within a single cave. The Warm River Cave stream is associated with an active hydrothermal system where we sampled the coldest accessible water. Cave access and sampling were conducted with appropriate permitting and permission from the respective landowners and managing agencies.

**Figure 1:**
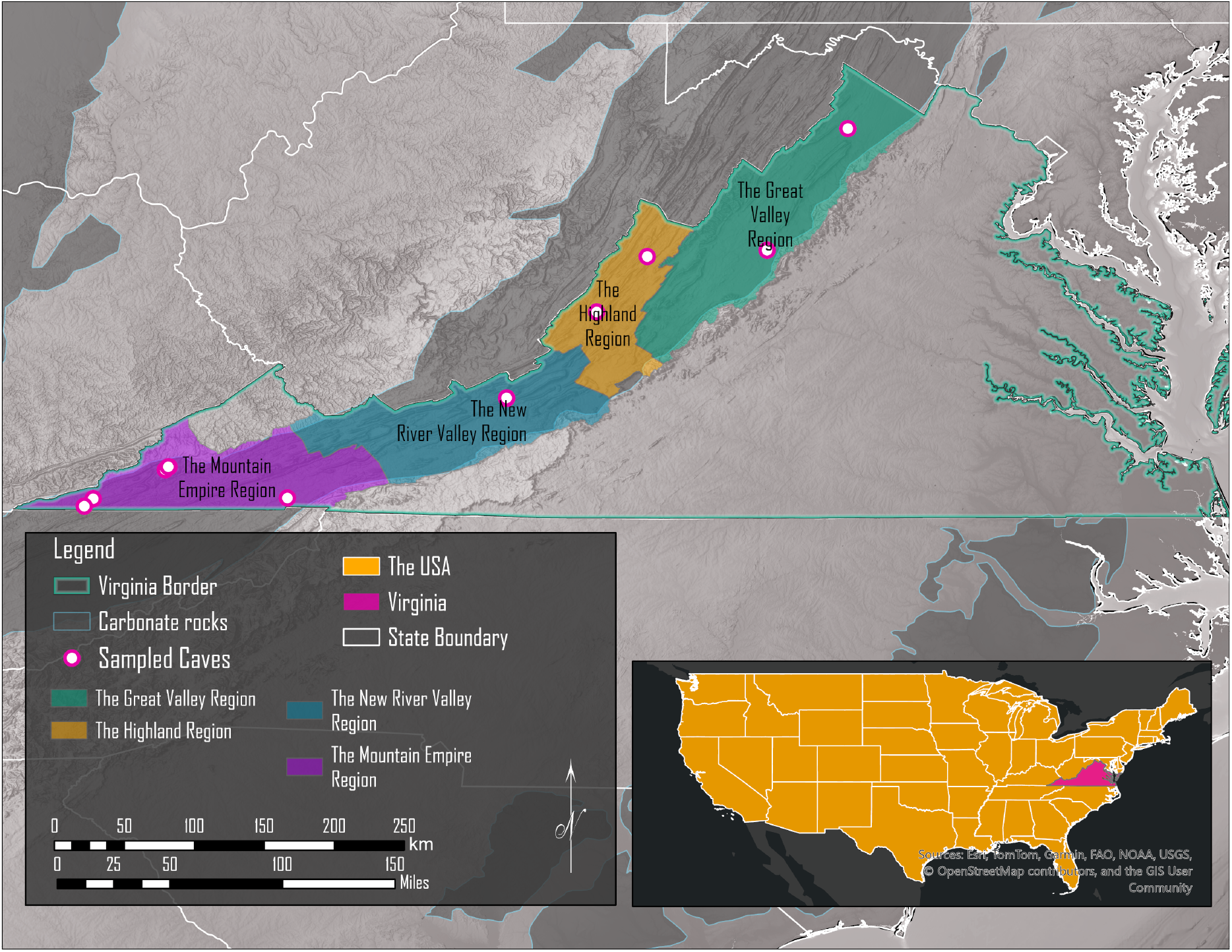
Cave water sampling sites distributed across the karst regions of Virginia, represented by white-filled magenta dots in color-shaded karst regions. Nine samples were collected from eight caves; the two Omega Cave samples (Omega Main Stream and Omega Tributary) share a single map location. Colored shading shows the major karst regions of Virginia. Cave names, counties, collection dates, overburden depth, host carbonate unit, information about position within the local flow system, and SRA accession numbers for each site are given in Table 3.

Air temperature and relative humidity (NIST-certified pocket meter, 32705K16; McMaster-Carr), pH, conductivity, salinity, and water temperature (PCTSTestr, 41115600; Oakton), and dissolved oxygen (EO600, 13945T62; Extech) were measured *in situ* and reported as metadata. Meters were calibrated daily before sampling against certified standards (pH 4.0, 7.0, and 10.0: 3108T35, 3108T25, and 3108T45; conductivity 84, 1,413, and 12,880 *µ*S: 3108T55, 3108T56, and 3108T65; all McMaster-Carr), and were calibrated at the sample site itself when the surface temperature differed from that of the site by more than 7 *^◦^*C. Readings were taken in a fixed order (air temperature, humidity, pH, conductivity, salinity, dissolved oxygen, water temperature) once instruments had equilibrated to the cave environment, defined as successive temperature readings within 0.2 *^◦^*C of one another.

### Field Procedures

Sampling site coordinates were recorded in reference to the nearest cave survey station, or, where no station was available, the distance, azimuth, and inclination from the sample site to a prominent landmark measured with a digital survey instrument (BRIC4; BricSurvey).

Samples were collected into 1 L gamma-sterilized bottles using a peristaltic pump with 6.4 mm (1/4 in.) inner-diameter Tygon tubing (XX800T25; Millipore), a barbed inline filter (8991T32; McMaster-Carr) to prevent large particles from being taken up in the sample, and a miniature suction strainer (9877K889; McMaster-Carr) to prevent macroscopic invertebrates from being drawn into the intake. The pump line was flushed with 80% bleach and then rinsed three times with sterile water immediately before sample collection; the third rinse was collected in a sterile bottle and retained as a process control. One process-control rinse control was collected per sample. Samples were transported to the lab in an insulated cooler with ice packs.

### Lab Procedures

A measured volume of 1L was vacuum-filtered onto two 0.22 *µ*m membrane filters (Advantec MFS, Inc) (22), for all samples, the full sample was passed– filter-clogging particles were prevented from being interoduced through the combination of basket and in-line filter. In cases where they could not be immediately processed, filters stored long term at *−*80 *^◦^*C prior to DNA extraction. DNA was extracted with the DNeasy PowerSoil Pro Kit (Qiagen). Full-length (V1–V9) 16S rRNA amplicons were generated by PCR with primers 27F (AGRGTTYGATYMTGGCTCAG) and 1492R (RGYTACCTTGTTACGACTT) (21). Amplicons were cleaned with AMPure XP beads (Beckman Coulter), and libraries were prepared with the SMRTbell Prep Kit (Pacific Biosciences) and sequenced on a PacBio Revio (25M SMRT Cell) using HiFi CCS (23).

### Computational Analysis

Reads were processed in R (24, 25) with DADA2 (26). Quality filtering retained reads of 1,350–1,650 bp containing no ambiguous bases (maxN = 0) with truncQ = 2 and no expected-error ceiling (maxEE = *∞*); PhiX-matching reads were discarded. Error models were learned separately for each flow cell, with a pooled model across all samples used as a fallback where per-flow-cell estimation failed to converge. Dereplicated reads were denoised with parameters appropriate for full-length CCS reads (band size = 32, OMEGA_A = 1*×*10*^−^*^40^). Amplicon sequence variants falling outside the 1,350–1,650 bp window were removed, and chimeras were then removed using the consensus de novo method implemented in DADA2 (removeBimeraDenovo) (26). ASVs were classified against SILVA v138.1 (27–29) at a minimum bootstrap confidence of 50, with species-level assignments added by exact matching. Contaminant ASVs were identified with decontam (30) using the prevalence method (threshold = 0.1, normalize = TRUE) with field rinse controls and extraction blanks as the negative control set; control samples were excluded from all downstream analyses. Mitochondrial and chloroplast sequences were also removed. This DADA2 workflow was used solely to enumerate amplicon sequence variants; ASV counts per sample are reported in Table 3. Due to the poor species-level resolution, no diversity indices were calculated from the ASV table.

All alpha- and beta-diversity statistics reported here were instead calculated with the EzBioCloud 16S-based Microbiome Taxonomic Profiling (MTP) pipeline (CJ Bioscience, Inc.; https://www.ezbiocloud.net) (31, 32), using the PKSSU5.0 reference database (analysis performed August 1st, 2026). Quality-controlled reads were assigned taxonomically against the EzBioCloud 16S database using VSEARCH (33), and operational taxonomic units (OTUs) were picked by the open-reference method: reads matching a reference sequence at *≥*97 % similarity were assigned to that species, unmatched reads were clustered de novo by UCLUST at a 97 % similarity boundary, and singleton de novo clusters were discarded. Alpha-diversity indices (, beta-diversity indices, and UPGMA clusering Samples were normalized to a common sequencing depth of 9,857 reads for alpha-diversity comparisons, full read counts were retained for differential-abundance testing (34, 35).

## Results

Sequencing yielded 205,617 raw reads across the nine samples (median, 13,551 reads/sample; range, 11,783–60,149), with a mean read length of 1,466.7 *±* 27 a mean Q score of 39.92 *±* 0.02, and GC content ranging from 52.42% to 54.75%. After processing, 3,899 unique ASVs were identified (recovered range 75–1,993 per sample), and these DADA2 processed reads were assigned to 37 phyla, 185 orders, 203 families, 355 genera, and 31 species.

To better understand the species-level diversity, the filtered reads were alternatively analyzed as OTUs using the PKSSU5.0 reference database. The family (Fig. 2) and species level diversity (Fig. **??**) are shown with the full complement of the reads passing filter. Alpha-diversity analyses were conducted following rarefaction to 9,857 reads (the lowest number of reads passing filter for this sample set) (Fig. 4. Shannon diversity ranged from 3.44 to 5.27 and Chao1 richness ranged from 426.11 to 1,896.96, indicating diverse assemblages that varied substantially among sites.

**Figure 2:**
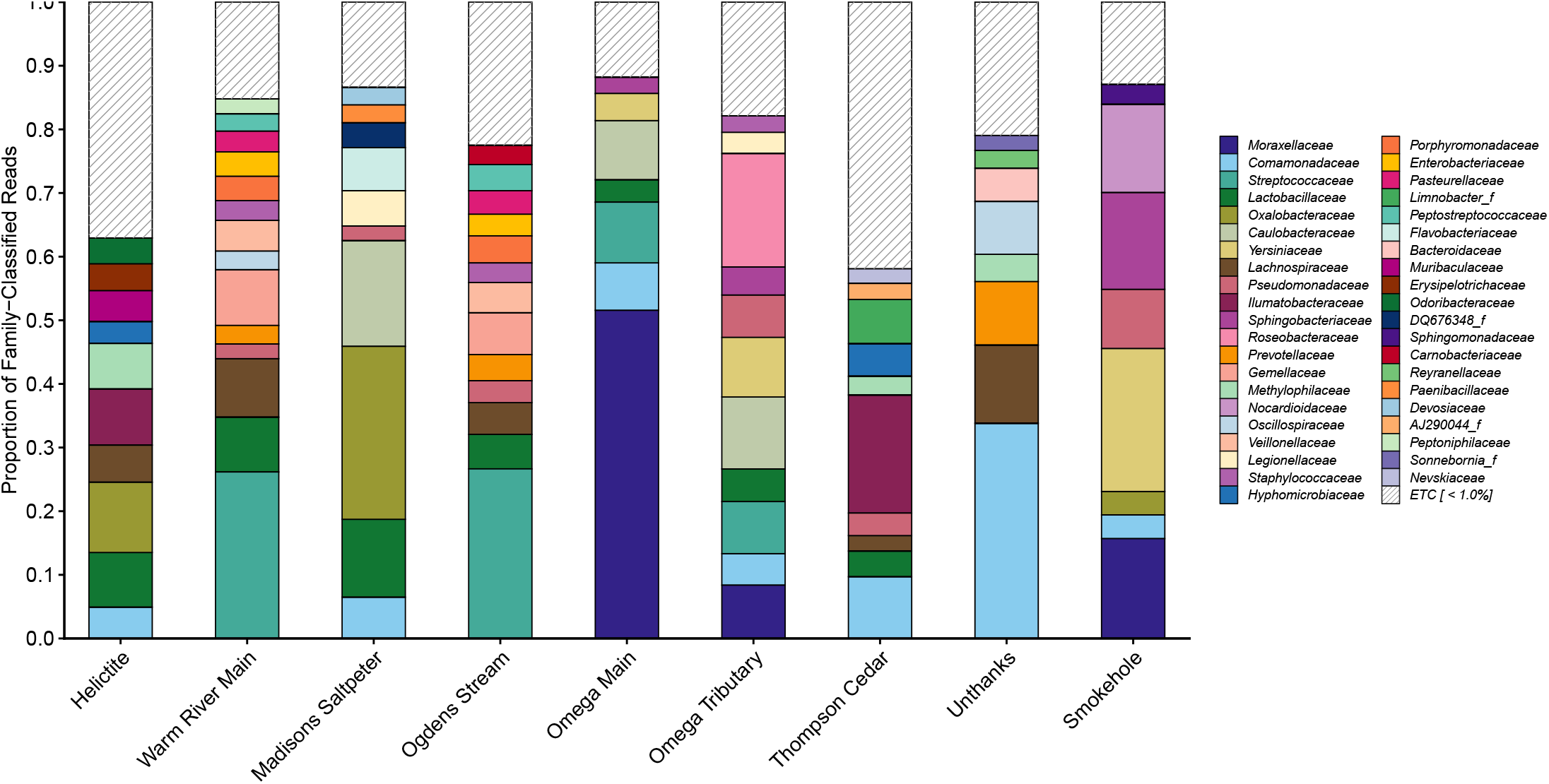
Relative frequency of families among family-classified reads across sites. Families that do not reach 1% of family-classified reads in at least one sample are pooled into the hatched category ETC [*<* 1.0%]; the same set of families is displayed for every sample. Reads that could not be assigned at the family level are excluded from the denominator, so bar heights describe composition within the classified fraction of each sample. Sites are ordered as in Table 3, by geologic region.

**Figure 3:**
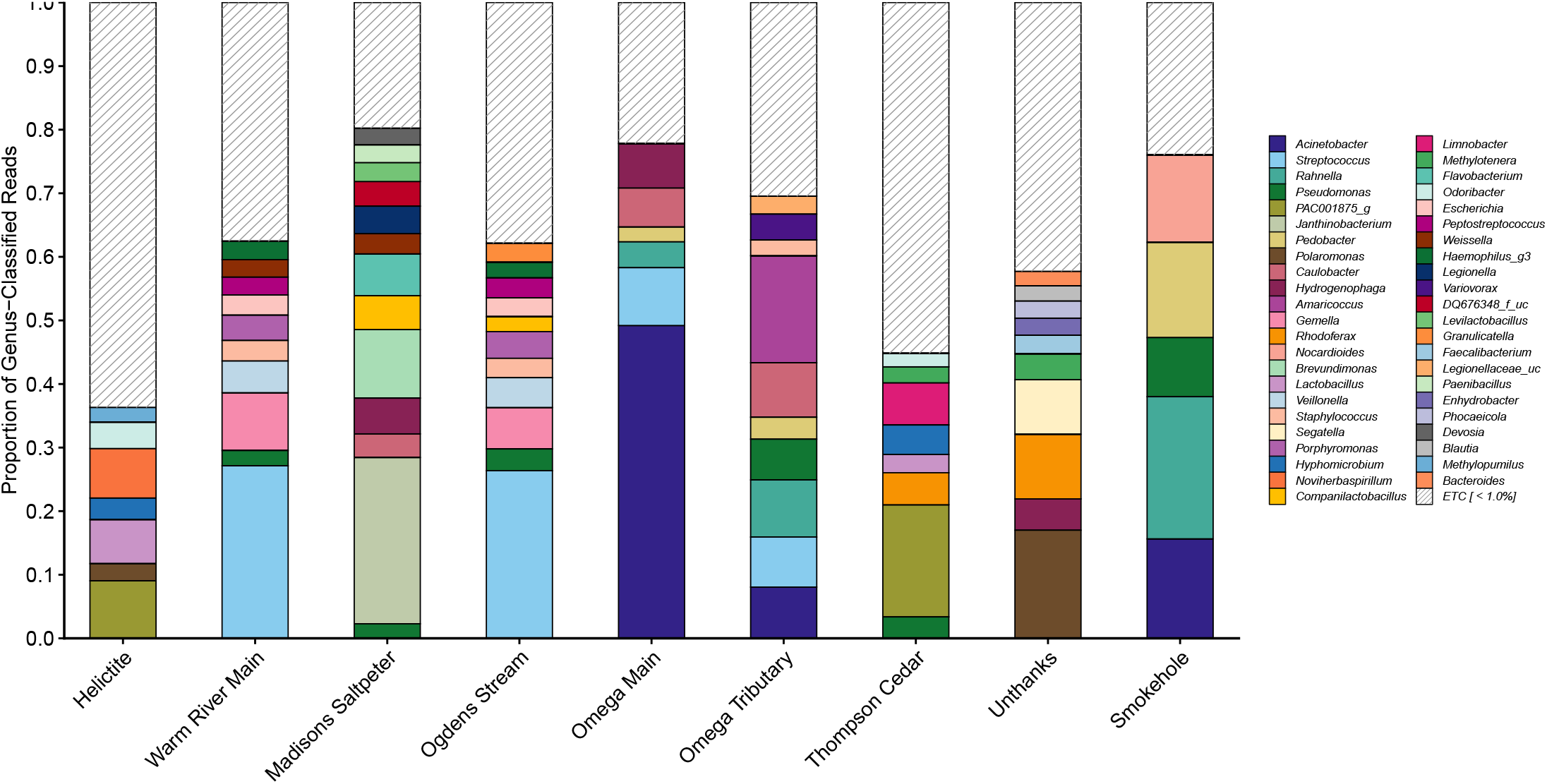
Relative frequency of genera among genus-classified reads across sites. Genera that do not reach 1% of genus-classified reads in at least one sample are pooled into the hatched category ETC [*<* 1.0%]; the same set of genera is displayed for every sample. Reads that could not be assigned at the genus level are excluded from the denominator, so bar heights describe composition within the classified fraction of each sample. Sites are ordered as in Table 3, by geologic region.

**Figure 4:**
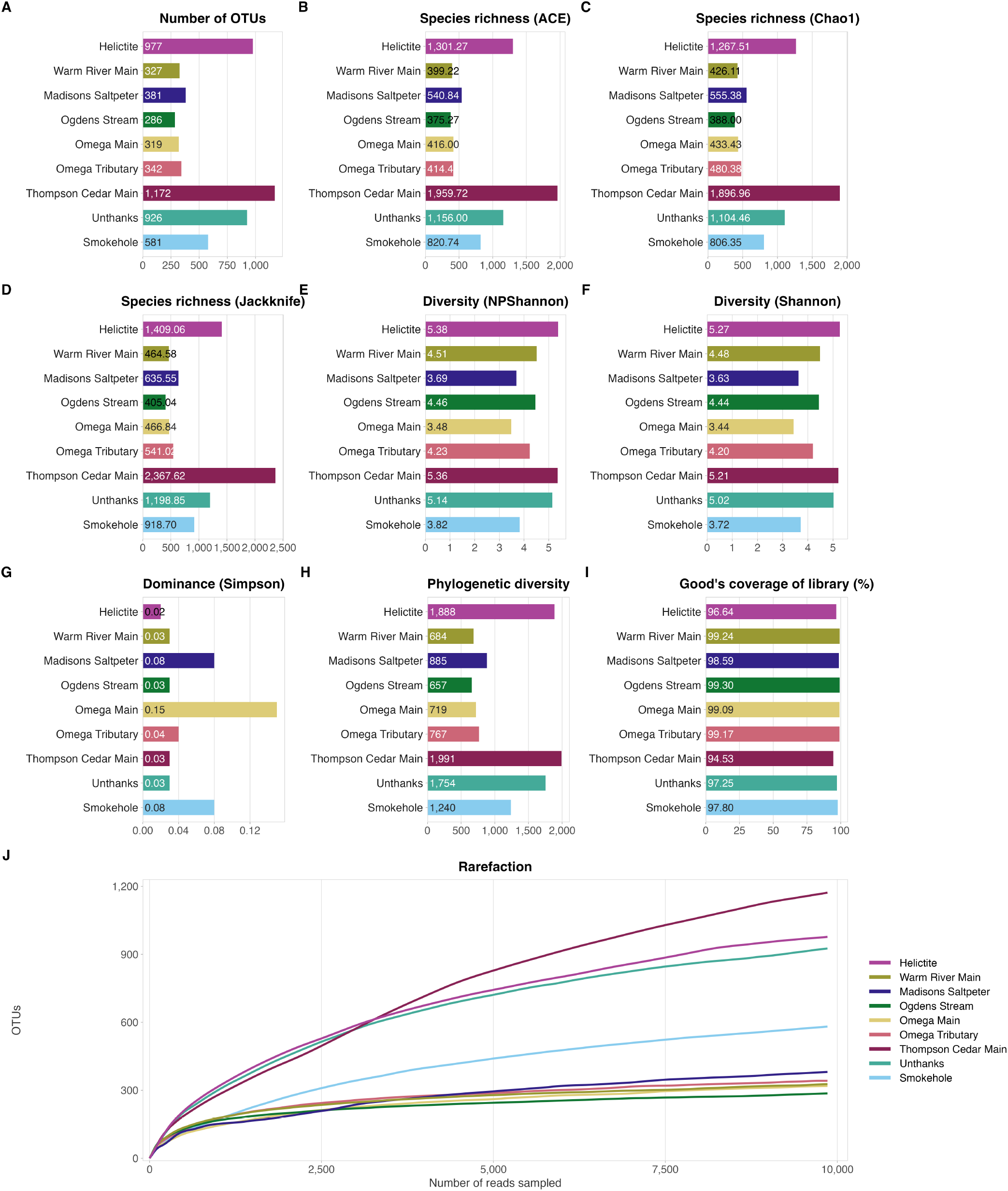
Alpha diversity of bacterial communities in nine Virginia karst groundwater sites. Operational taxonomic units (OTUs) were assigned by open-reference UCLUST clustering at a 97% sequence identity threshold (minimum cluster size 2) against the ChunLab reference database, restricted to bacterial 16S rRNA gene sequences. To permit comparison across samples of unequal sequencing effort, all libraries were normalised by random subsampling to a common depth of 9,857 reads per sample; every index reported here is calculated at that depth. (A) Observed OTU richness. (B–D) Estimated richness by the ACE, Chao1 and jackknife estimators. (E, F) Shannon diversity, calculated by the non-parametric (NPShannon) and classical estimators respectively. (G) Simpson dominance index, for which larger values indicate stronger dominance by a small number of taxa. (H) Faith’s phylogenetic diversity. (I) Good’s coverage of the library, expressed as a percentage. In panels A–I, bars are ordered from top to bottom by karst region and the numerical value of each index is printed on each bar. (J) Rarefaction of OTU richness against sequencing depth. Rarefaction curves in panel J were computed by subsampling without replacement and are shown to the 9,857-read normalisation depth.

UPGMA clustering recovered the same closest pair under all four distance metrics examined and at both taxonomic levels (Fig. 5): Ogdens Stream and Warm River Main joined at the shortest distance under Jensen–Shannon, Bray–Curtis, generalized UniFrac, and UniFrac distances computed at genus and at species level.

**Figure 5:**
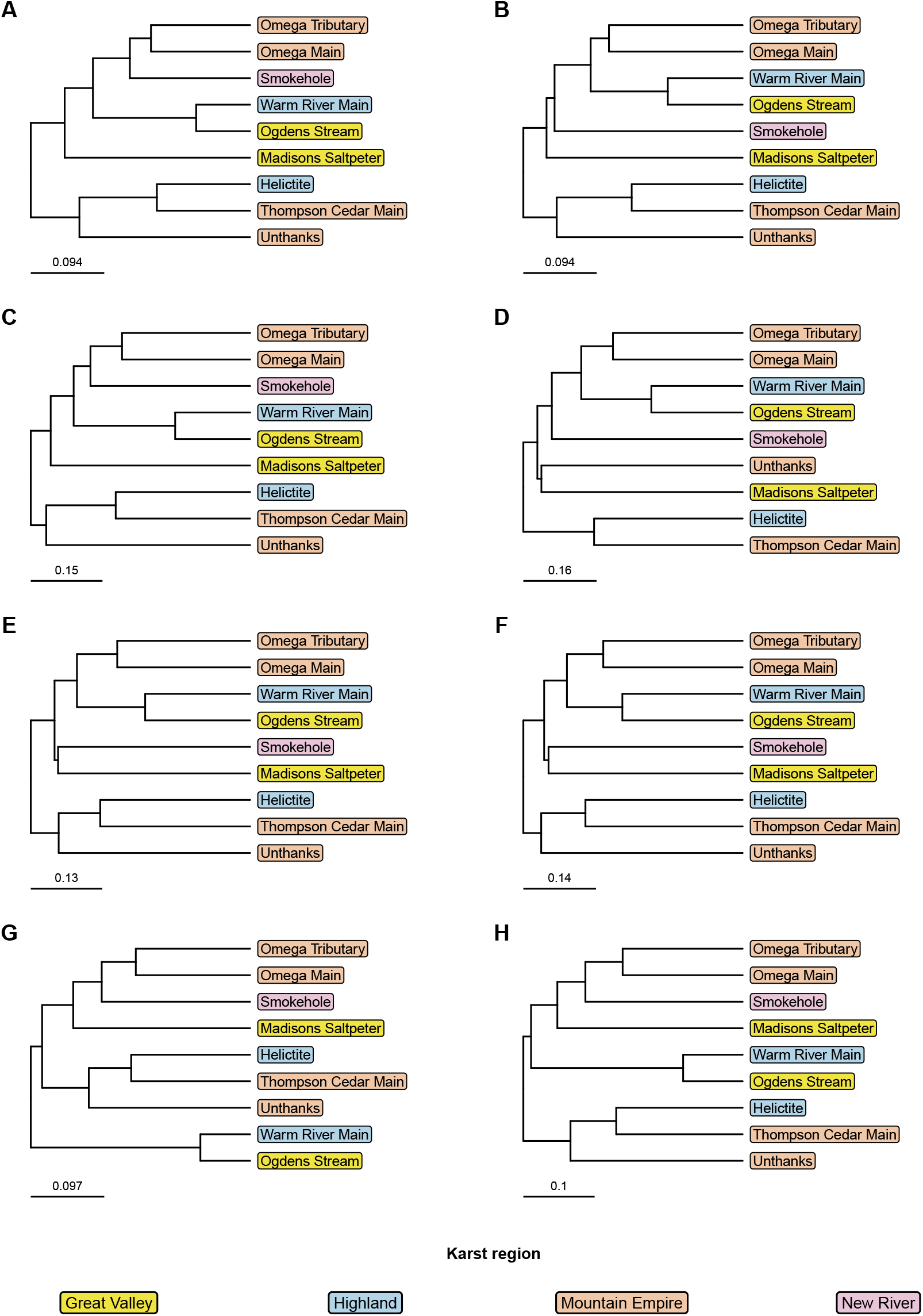
UPGMA clustering of beta-diversity distances among nine Virginia karst groundwater microbial communities. Dendrograms were generated by unweighted pair group method with arithmetic mean (UPGMA) clustering of pairwise beta-diversity distances calculated in the EzBioCloud MTP Set Browser (ChunLab, Seoul, Republic of Korea) from 16S rRNA gene amplicon profiles, using database version PKSSU5.0. (A) Jensen–Shannon divergence computed at the genus level. (B) Jensen–Shannon divergence computed at the species level. (C) Bray–Curtis dissimilarity computed at the genus level. (D) Bray–Curtis dissimilarity computed at the species level. (E) Generalized UniFrac distance computed at the genus level. (F) Generalized UniFrac distance computed at the species level. (G) UniFrac distance computed at the genus level. (H) UniFrac distance computed at the species level. Distances were calculated without normalization, and reads assigned to unclassified OTUs were excluded. Scale bars indicate distance in the units of the metric shown in that panel; scales are not comparable across panels.

In a principal coordinates analysis of genus-level Jensen–Shannon distances (Fig. 6), the first two axes accounted for 45.8% and 20.8% of the total variation (66.6% cumulative), with a third axis, not shown, explaining a further 14.1%.

**Figure 6:**
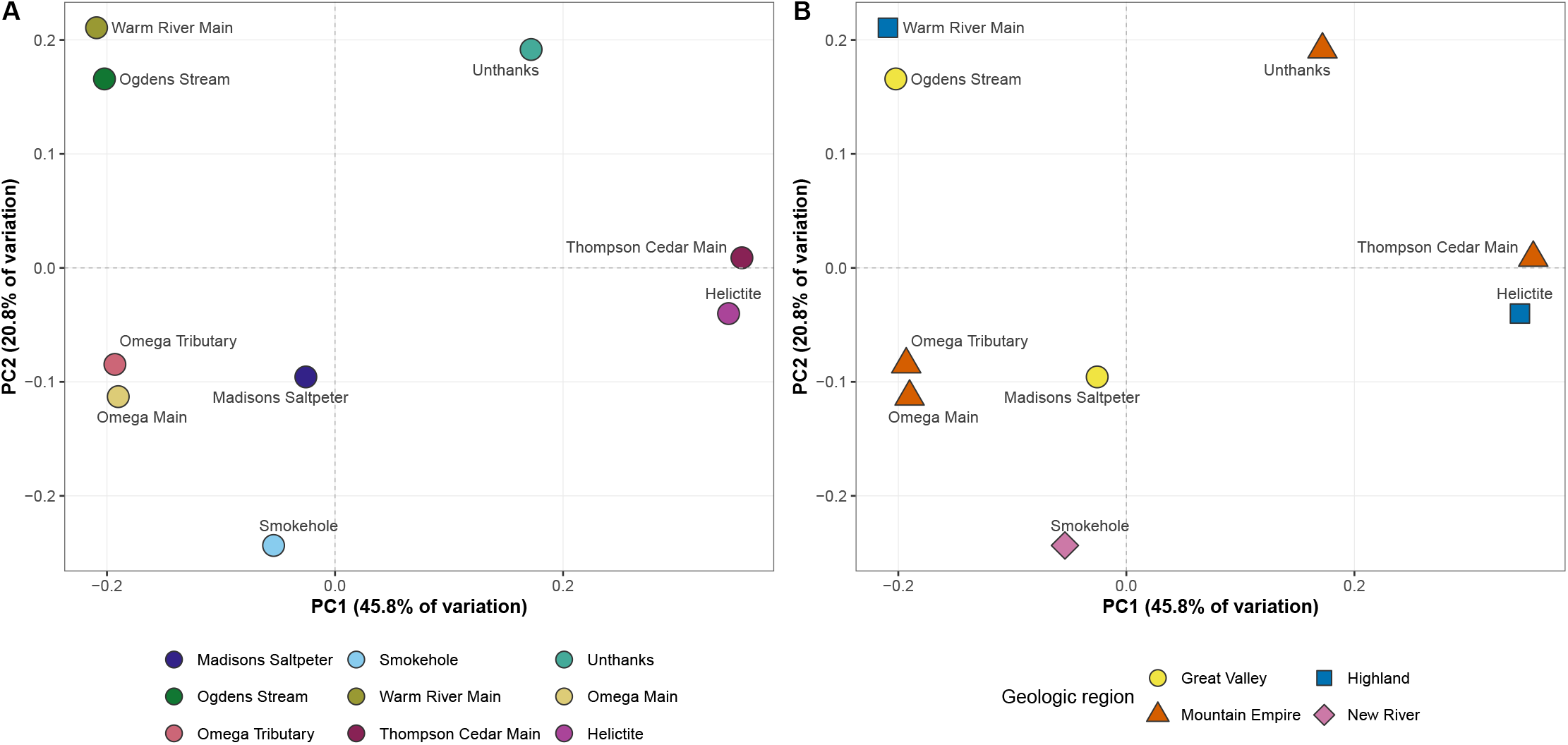
Beta diversity of bacterial communities in nine Virginia karst groundwater sites. Ordination is based on Jensen–Shannon distances computed from genus-level community profiles, with reads from unclassified OTUs excluded; libraries were normalised to 9,857 reads per sample prior to distance calculation. Taxonomic assignment was performed against the EzBioCloud PKSSU5.0 16S rRNA database. The first two principal coordinates explain 45.8% and 20.8% of the total variation respectively (66.6% cumulative); a third axis, not shown, explains a further 14.1%. Each point represents one sample, coloured by site; axes are drawn to a common scale so that inter-point distances are directly comparable.

Reads were also assigned to likely metabolic guilds on the basis of genus-level taxonomy (Fig. 7). All eleven guilds were represented in at least one site. The microbial assemblages were dominated by host-associated lineages, followed by aerobic copiotrophs, and aerobic oligotrophs. Host-associated taxa were recovered from all nine sites but at markedly different levels (Table 2). The assemblage associated with the gastrointestinal tract of humans and livestock accounted for 58.5% of genus-level reads at Warm River Main and 57.4% at Ogdens Stream, 17.5% and 17.0% at Omega Main and Omega Tributary, and 15.5% at Unthanks, but no more than 3.9% at the four remaining sites. A distinct wildlife-associated gut assemblage showed the opposite pattern, reaching 12.5% at Helictite and 5.8% at Thompson Cedar Main, the two sites at which the human/livestock assemblage was among the lowest recorded (2.0% and 0.8% respectively). Process controls carried alongside the samples yielded between four and ten sequences each.

**Figure 7:**
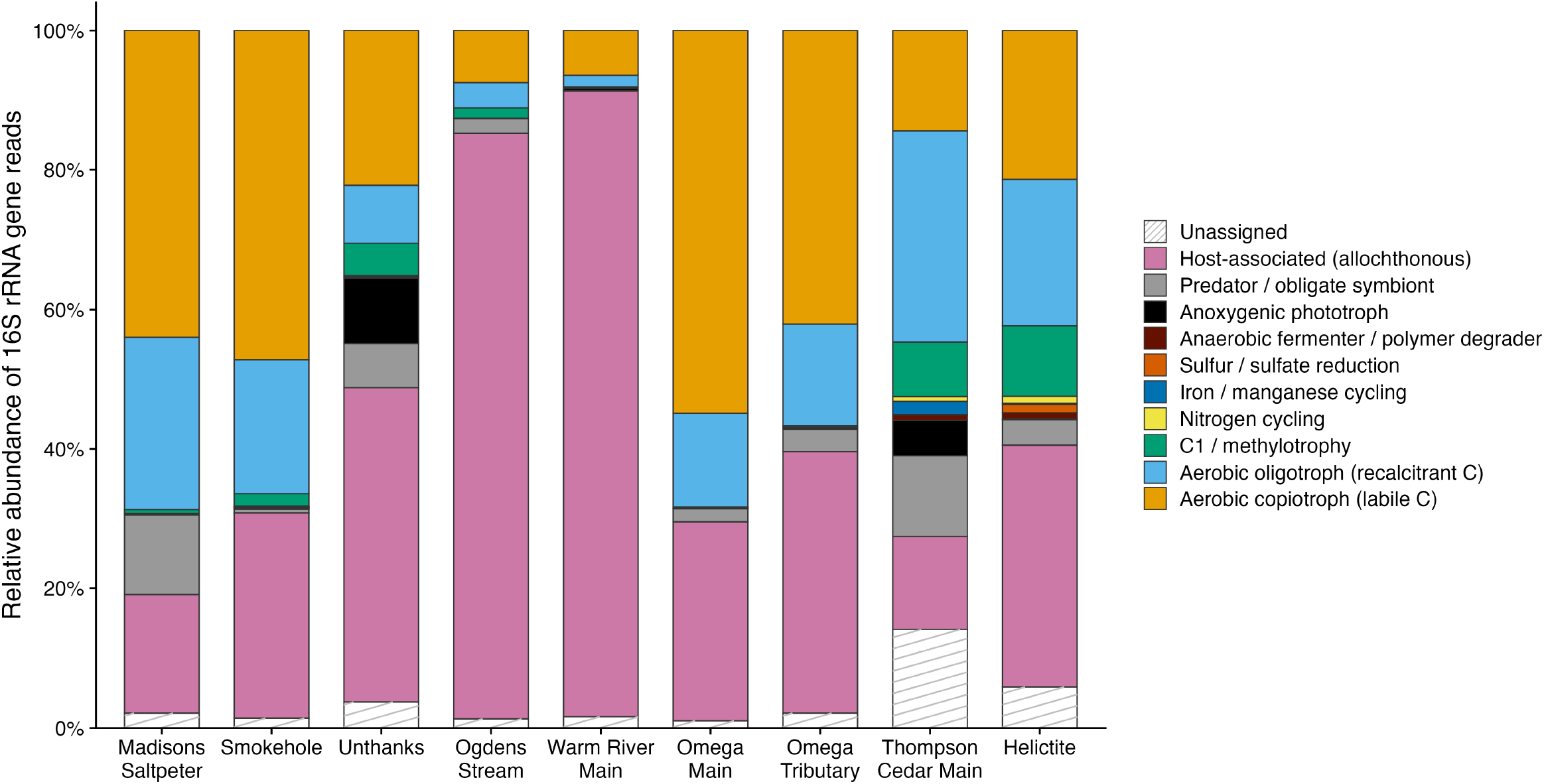
Metabolic guild composition of bacterial communities in nine Virginia karst groundwater sites. Genus-level 16S rRNA gene reads were assigned to eleven functional guilds on the basis of taxonomic affiliation, with each genus allocated to a single guild. Bars give the relative abundance of reads per guild within each library; all eleven guilds are represented in at least one site. The *Unassigned* category is drawn in white with grey hatching rather than as a further color, to mark it as differing in kind from the guilds: it records taxa for which the reference taxonomy supports no functional inference, not an inferred strategy. Guild assignment is inferred from taxonomy and is not a measurement of activity; it should be read as a description of the metabolic strategies the resident lineages are capable of, not as a precise description of the metabolic processes operating at the time of sampling.

**Figure 8:**
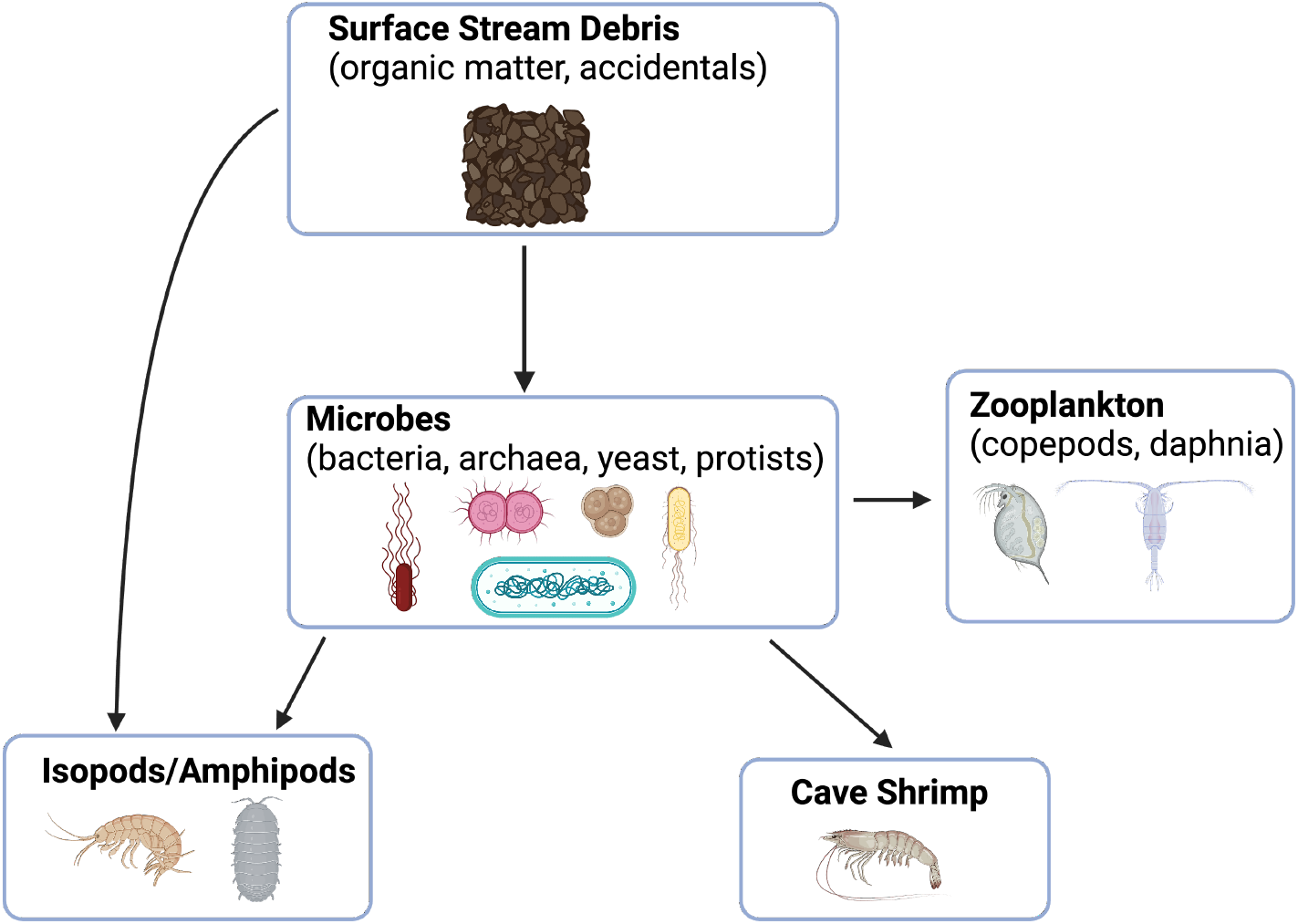
Hypothesized food web diagram for aquatic cave habitats. Created with BioRender. Adapted from Helf and Olson (52); original figure modified from Barr and Kuehne (53).

**Table 1:** Taxonomy richness by OTUs, before subsampling (full cleaned data). Values are the number of distinct taxa with at least one read assigned in each sample; the “Unclassified in higher taxonomic rank” bin is excluded. The “Observed” row gives the number of taxa observed in at least one sample, not the column sum.

| Sample | Phylum | Class | Order | Family | Genus | Species |
| --- | --- | --- | --- | --- | --- | --- |
| Helictite | 37 | 88 | 184 | 306 | 664 | 1214 |
| Madisons Saltpeter | 23 | 44 | 87 | 138 | 236 | 375 |
| Ogdens Stream | 14 | 27 | 61 | 98 | 181 | 288 |
| Omega Main | 17 | 35 | 71 | 110 | 204 | 343 |
| Omega Tributary | 16 | 31 | 72 | 114 | 210 | 347 |
| Smokehole | 24 | 48 | 114 | 172 | 343 | 595 |
| Thompson Cedar Main | 44 | 107 | 227 | 401 | 799 | 1300 |
| Unthanks | 30 | 67 | 144 | 225 | 489 | 841 |
| Warm River Main | 9 | 19 | 53 | 87 | 190 | 325 |
| Observed (all samples) | 54 | 135 | 296 | 569 | 1493 | 3350 |

**Table 2:** Relative abundance (% of genus-level reads) of host-associated assemblages at each site, with geologic region and county human population density. Assemblage membership was assigned at genus level. Genera associated with the oral cavity and gastrointestinal tract of humans and livestock are reported as a single group, as many of the taxa concerned occur in both compartments and are not resolved to host compartment by genus assignment alone. Rows are not additive, as a small number of genera occur in more than one assemblage. Population densities are from the 2020 U.S. Decennial Census, the census nearest the 2022 sampling campaign, and are reported for the county containing each site. Virginia’s independent cities are separate jurisdictions and are excluded from the surrounding county’s figure; see footnote. Sites are listed in the order of Table 3.

| Cave | Geologic region | County | Population density (persons km<sup>-2</sup>) | Valid reads (analyzed) | Human/livestock GI tract | Wildlife GI tract |
| --- | --- | --- | --- | --- | --- | --- |
| Helictite | Highland | Highland | 2.1 | 51,514 | 2.0 | 12.5 |
| Warm River Main | Highland | Alleghany | 13.2 | 9,857 | 58.5 | 2.5 |
| Madisons Salt-peter | Great Valley | Augusta | 30.9<sup>a</sup> | 10,999 | 3.1 | 0.1 |
| Ogdens Stream | Great Valley | Frederick | 85.4<sup>b</sup> | 11,331 | 57.4 | 1.1 |
| Omega Main | Mountain Empire | Wise | 34.6<sup>c</sup> | 12,408 | 17.5 | 1.3 |
| Omega Tributary | Mountain Empire | Wise | 34.6<sup>c</sup> | 11,863 | 17.0 | 1.9 |
| Thompson Cedar Main | Mountain Empire | Lee | 19.7 | 56,364 | 0.8 | 5.8 |
| Unthanks | Mountain Empire | Lee | 19.7 | 11,361 | 15.5 | 0.3 |
| Smokehole | New River | Giles | 18.1 | 12,098 | 3.9 | 0.7 |

<sup>a</sup>Excludes the independent cities of Staunton (499 persons km<sup>-2</sup>) and Waynesboro (572 persons km<sup>-2</sup>), which are surrounded by but not part of Augusta County.
<sup>b</sup>Excludes the independent city of Winchester (1,181 persons km<sup>-2</sup>), which is surrounded by but not part of Frederick County.
<sup>c</sup>Excludes the independent city of Norton (190 persons km<sup>-2</sup>), which is surrounded by but not part of Wise County.

**Table 3:** Description of sample sites and accession numbers (SRA) for each.

| Cave | Geologic region | County | Date of sample collection | Depth of collection site, overburden (ft) | Carbonate unit | Closer to input or discharge of cave system? | Total reads (raw) | Valid reads (analyzed) | ASVs inferred | SRA |
| --- | --- | --- | --- | --- | --- | --- | --- | --- | --- | --- |
| Helictite | Highland | Highland | 25 April 2022 | 236 | Helderberg Group (Siluro-Devonian) | Input | 54,701 | 51,514 | 1993 | SRR37725181 |
| Warm River Main | Highland | Alleghany | 26 April 2022 | 317 | Middle Ordovician | Discharge | 11,783 | 9,857 | 83 | SRR37725173 |
| Madisons Salt-peter | Great Valley | Augusta | 23 April 2022 | 171 | Conococheague Formation (Cambrian-Ordovician) | Phreatic Zone | 11,914 | 10,999 | 85 | SRR37725180 |
| Ogdens Stream | Great Valley | Frederick | 24 April 2022 | 25 | Beekmantown Formation (Ordovician) | Input | 13,561 | 11,331 | 104 | SRR37725179 |
| Omega Main | Mountain Empire | Wise | 26 November 2022 | 540 | Greenbrier (Mississippian) | Input | 13,904 | 12,408 | 187 | SRR37725178 |
| Omega Tributary | Mountain Empire | Wise | 26 November 2022 | 496 | Greenbrier (Mississippian) | Input | 13,082 | 11,863 | 267 | SRR37725177 |
| Thompson Cedar Main | Mountain Empire | Lee | 28 April 2022 | 268 | Hurricane Bridge (Ordovician) | Discharge | 60,149 | 56,364 | 1205 | SRR37725175 |
| Unthanks | Mountain Empire | Lee | 28 April 2022 | 90 | Martin Creek (Ordovician) | Discharge | 13,551 | 11,361 | 75 | SRR37725174 |
| Smokehole | New River | Giles | 1 May 2022 | 70 | Middle Ordovician | Discharge | 12,972 | 12,098 | 269 | SRR37725176 |

## Discussion

These data provide what is, to our knowledge, the first survey of the aquatic microbiota of cave waters across Virginia and reveal diverse assemblages that differed markedly among the sampled caves. The Warm River Cave stream sample was collected from a hydrothermal cave system. The pronounced differences in composition and diversity among sampled sites may indicate that local conditions—hydrology, geology, and organic-matter inputs—shape these communities. Our current dataset is too limited to understand the significance of geochemical or environmental variables driving compositional differences, but continued sample collection paired with collection of geochemical and environmental variables may reveal parameters that most strongly influence microbial community composition in this setting. We expect, for example, that Total Organic Carbon and dissolved oxygen will play a role in nutrient availability.

We propose that sequencing microbes in cave waters could be a particularly promising method for rapid, cost-effective, and sensitive environmental monitoring (36–38). Microbes are particularly effective indicators of environmental changes because they are both metabolically specialized and fast-multiplying (37, 39). When the types of nutrients available in an environment change, the population of microbes that lives on the dominant nutrient sources will quickly expand, while the population that relied on the now-absent nutrients will rapidly contract (40, 41). Baseline measurements of microbial communities are the most rigorous way to understand changes in microbial communities resulting from changing nutrient availability (38, 42), but in the case of known surface contamination events, such as an oil spill, microbial profiling can help land managers see how quickly microbes, which serve as the base of the cave ecosystem (43, 44), are responding to environmental changes (45). Figure 7 shows the metabolic potential of the sampled microbial community, as predicted by genus-level assignments of DNA profiles (39).

While data about metabolic strategies in immediate use by the microbial community would require RNA profiling (46), karst systems offer an unusually direct route for a surface contaminant to reach the subsurface, and bacterial assemblages in karst springs and epiphreatic pools shift measurably over the course of individual recharge events (47–49). We therefore expect that a contamination event would register in the community profile on the timescale of days, though we note that such a shift may reflect the import of surface-derived taxa as much as *in situ* growth of subsurface degraders (48, 50), and that karst groundwater communities are otherwise taxonomically stable relative to the nutrient-replete surface waters in which rapid substrate-driven succession has been most fully characterised (40, 47, 51).

## Caveats and Limitations

### Predicted Provenance of Host-Associated Bacteria

These samples were taken across diverse karst areas, whose watersheds included both urban and agricultural land areas. It is thus not surprising that host-associated (mammalian) taxa were recovered from all nine sites. These host-associated bacteria can be separated by probable source, and each of these groups has a different site distribution.

The first group is an assemblage of taxa associated with the oral cavity and gastrointestinal tract of humans and livestock, which dominated Ogdens Stream and Warm River Main and was substantial at the two Omega sites. *Streptococcus* alone accounted for 24.2% and 24.4% of reads at Ogdens Stream and Warm River Main respectively, and was accompanied by the genera with which it co-occurs on mucosal surfaces—*Gemella* (6.0%, 8.2%), *Veillonella* (4.3%, 4.5%), *Porphyromonas* (3.9%, 3.6%), *Granulicatella*, *Haemophilus*, *Neisseria*, *Rothia*, *Fusobacterium*, *Capnocytophaga*, and *Abiotrophia* (54). *Streptococcus*, *Veillonella*, *Fusobacterium*, *Gemella*, *Granulicatella*, *Rothia* and *Neisseria* have all been recovered from the rumen and lower gastrointestinal tract of cattle, sheep and other ruminants (55, 56). The same assemblage includes the obligately anaerobic gut taxa that peaked at Unthanks (12.9% of reads), where *Faecalibacterium* (2.7%), Blautia (2.2%), B*acteroides* (2.1%), *Agathobacter* (1.0%), *Akkermansia* (1.0%), and *Roseburia* (0.9%) were all present, with Lachnospiraceae and Oscillospiraceae at 10.8% and 7.3% at family level. Summed across both mucosal and anaerobic gut members, the assemblage reached 57.4% and 58.5% at Ogdens Stream and Warm River Main, 15.5% at Unthanks, and 17.5% and 17.0% at Omega Main and Omega Tributary, against *≤*3.9% at the remaining four sites (Table 2).

The second host-associated group, a gut assemblage more typical of wildlife, characterized Thompson Cedar Main and Helictite: Muribaculaceae (1.9% and 4.5%), *Odoribacter* (1.9% and 3.7%), Rikenellaceae, *Turicibacter*, and *Lactobacillus* (2.6% and 6.3%). Muribaculaceae is a host-restricted family typical of rodent and other small-mammal guts rather than of human or livestock faeces (57), and the genera of the first assemblage were essentially absent from these two sites (*≤*0.6% and *≤*1.4% respectively). Resident cave fauna, most plausibly bats or small mammals, are the likely importers of these microbial communities.

Process controls carried alongside the samples yielded between four and ten sequences each, a level indistinguishable from index bleed-through and far below any of the abundances reported here. Reagent-and kit-derived contamination, to which low-biomass groundwater samples are otherwise highly susceptible (58–62), can therefore be excluded as the origin of these assemblages.

### Comparisons & Future Directions

Comparable host-associated signals have been reported across a range of karst settings and land-use intensities. In the urban karst of Bowling Green, Kentucky, clinically relevant resistance determinants were recovered year-round from springs, sinkholes, and wells irrespective of karst feature type or season (22, 63). Paired urban and peri-urban sites in Chongqing, China differed in both community composition and resistome, with the more heavily developed catchment enriched in human-associated taxa (64). Across eight Romanian karst springs, low-abundance host-associated lineages tracked surface inputs which introduced variability to an otherwise stable core community (65). At the opposite end of the gradient, some alpine karst aquifers that lack anthropogenic impacts have been described as dominated by a taxonomically stable autochthonous community in which host-associated taxa appear only transiently and in association with high-discharge events (66, 67). For example, in a Sicilian cave stream a single runoff event was sufficient to displace the resident sulfur-oxidizing community with surface- and anthropogenically derived taxa (68). The Virginia caves sampled here are somewhere between the urban karst of Kentucky and undisturbed alpine karst sites like those sampled in Romania– most of the caves we sampled were in rural and agricultural settings, located below land where people live and work.

Paired sampling of surface recharge points and cave streams, host-specific *Bacteroidales* source-tracking markers, and quantification of DNA yield per litre at multiple points along the flow path would test the recharge pathway directly. Further, sampling across a hydrograph rather than at base flow alone would distinguish transient recharge-associated microbial inputs from a persistent resident microbial community (67, 68).

### Limitations of this Study

An important caveat is that the sampled waters are cave waters and therefore represent only a subset of the broader karst aquifer system. Cave passages afford access to groundwater at selected points along a flow path. Most karst water used by people is withdrawn from wells or discharged at springs rather than from the cave streams and pools sampled in this study, and the microbial communities at those points of use may or may not resemble those observed in cave waters. A direct comparison of cave, spring, and well microbiomes within the same flow systems would be a valuable next step and would clarify how representative cave-water communities are of the aquifer as a whole.

Sampling was not distributed evenly through the year: seven samples were collected in late April- early May 2022 and two (both from Omega Cave) in late November 2022. Season is therefore partially confounded with site.

Our findings are limited by the small number of samples and single-timepoint sampling. Because conventional inventories of cave fauna generally require repeated visits before the full diversity present can be reliably estimated (12–16), we propose that the single-visit sampling like that reported here is best used as a complement to a biological inventory and is not meant to replace a complete inventory of these systems.

In general, we refrain from drawing robust conclusions about the relationships between microbial communities or their expected microbial occupants from a single observation. Some karst environments may be oligotrophic, and these environments can yield highly diverse assemblage, which when sequenced at moderate depth can yield poorly overlapping subsets of otherwise similar communities (69). Repeated sampling and deeper sequencing of these cave streams will be needed to establish whether the difference between these microbial communities is reproducible or an artifact of medium-depth sequencing. While rarefaction curves in Figure 4(J) cannot be directly compared to the accumulation curves used in invertebrate sampling (15), rarefaction curves can be used to estimate and bound the proportion of the microbial community sequenced in a given sample. As is the case with invertebrate sampling, some seasonal variation is expected in the aquatic microbial occupants of the cave as a result of changing nutrient influx, and this should be taken into account when calculating appropriate sequencing depth and number of sampling iterations required to characterize aquatic microbial communities.

Initial sequencing of microbial DNA directly from cave water can be accomplished in a single visit, and these results may help establish reference points for future work aimed at monitoring how seasonal cycles and surface disturbances affect microbial communities in these caves with implications for the management of municipal water supplies and for the conservation of the region’s approximately 200 endemic cave invertebrates.

## Data availability

Sequencing data and associated metadata generated by this study are publicly available in the National Center for Biotechnology Sequence Read Archive under BioProject accession number PRJNA1441205. The individual accession numbers (SRR37725173–SRR37725181) and hyperlinks are included in Table 3.

## Funding Sources

This work was supported by a grant from the Cave Conservancy of the Virginias and through assistance from the Virginia Department of Conservation and Recreation, Natural Heritage Program.

## Acknowledgements

We are grateful for the assistance of the following individuals who helped our team transport sampling equipment to and from remote locations and helped with sample site documentation: Laura Young, Caleb Haines, Jason Delafield, Yvonne Droms, Mike Ficco, Bill Koerschner, Eli Meyer, Mark Minton, and Dave Socky.

## Competing interests

The authors declare no competing interests.

## Notes

https://www.ncbi.nlm.nih.gov/bioproject/PRJNA1441205

### Competing Interest Statement

The authors have declared no competing interest.

